# Multimodal Alignment of MicroCT Imaging to Vibroacoustic Signals to Validate Soft Tissue Needle Transitions in *Manduca sexta*

**DOI:** 10.64898/2026.06.07.730726

**Authors:** K. Steeg, R. Urrutia, A. Illanes, P. Fuentealba, J. Gawron, K. Strama, C. Hansen, J. Scherberich, A. Windfelder, G. Krombach, M. Friebe

## Abstract

**Objective:** Robotic-assisted needle insertions lack haptic feedback, a key sensory cue for detecting tissue transitions and regulating puncture force. Modeling this feedback requires an understanding of soft-tissue biomechanics during insertion. Vibroacoustic signals generated by needle–tissue interactions may provide an additional sensing modality, but their interpretation requires validation against anatomical ground truth.

**Methods:** A multimodal framework was developed to correlate vibroacoustic signals with high-resolution post-puncture microCT (µCT) imaging in *Manduca sexta*, an insect model containing interconnected soft-tissue layers. A custom clip-on prototype recorded vibroacoustic signals during manual needle insertions. Three trajectory-marking strategies were evaluated to determine 3D coordinates of soft-tissue layer crossings and to assess correlations between acoustic events and anatomical transitions. Distances between layer crossings and needle displacement were used for spatiotemporal alignment of vibroacoustic and µCT data.

**Results:** A µCT-compatible nylon string preserved puncture trajectories without artifacts and enabled high-resolution 3D reconstruction of anatomy and needle paths. Fusion of vibroacoustic and imaging data allowed identification of acoustic events associated with tissue entry, exit, and transitions.

**Conclusion:** By integrating high-resolution µCT imaging with vibroacoustic sensing, this study establishes a biologically grounded framework for validating the relationship between vibroacoustic signals and anatomical tissue transitions during needle insertion, providing a basis for future quantitative analyses.

**Significance:** This work provides initial evidence for correlating vibroacoustic signals recorded during needle insertion with corresponding µCT-identified tissue barriers. Because vibroacoustics offers substantially higher temporal and spatial resolution than most imaging modalities, it has the potential to improve tissue sensing and procedural accuracy in future needle-based interventions.

## I. Introduction

Accurate tissue differentiation during needle placement is critical in many minimally invasive procedures, such as obtaining biopsy samples. In addition to imaging modalities, which often lack sufficient resolution, haptic feedback is a major source of information about tissue boundaries for the clinician [1], [2]. This feedback arises from biomechanical needle-tissue interactions, so-called friction-induced vibrations (FIV), which encode real-time information about anatomical boundaries and procedure progress [2], [3].

Similarly to manual insertions, haptic feedback is needed in robotic-assisted interventions for dynamic force regulation and intraoperative procedure assessment. But in contrast to a human, a machine does not “sense” this feedback but relies on mathematical simulations and sensors, limiting their clinical impact due to challenges in achieving high-sensitivity and sufficient precision [2]. Furthermore, simulations are based on simplified assumptions that do not fully represent the complexity of biological tissues [4], [5]. Multimodal fusion has therefore emerged to combine complementary data sources such as imaging, force, and acoustic sensing for a more comprehensive understanding of the puncture environment [2], [6].

Within this field, vibroacoustic sensing represents a novel modality that translates needle–tissue interactions into measurable signals, potentially offering a novel real-time feedback channel for manual and robotic procedures. From a physical perspective, vibroacoustic signals arise from intraoperatively generated FIV that propagate along the needle shaft. The shaft acts as an oscillating resonator transmitting these vibrations to a proximally placed sensor. [7], [8] A long-term objective of this technology is to achieve real-time tissue characterization based on vibroacoustic signatures. Several studies assessed the feasibility of distinguishing between different homogeneous materials and tissues using vibroacoustic signals obtained during palpation or puncture [9], [10]. However, translating this approach into clinical practice faces significant challenges because it requires extensive datasets that establish comprehensive signal libraries to account for the variability and heterogeneity of biological tissues.

Another objective utilizing vibroacoustic signals is the detection of discrete anatomical events, such as transitions between tissues or layer crossings. Studies on artificial phantoms showed that vibroacoustics carry structurally meaningful information and have indicated that even thin layers, such as paper sheets, placed between soft materials create detectable vibroacoustic events [11]. Additionally, vibroacoustic signals obtained from puncturing artificial soft tissues embedded in gelatine, enabled the distinction between the initial puncture event when entering the material and moving through tissue with 95 % accuracy [12]. Furthermore, studies in *ex vivo* tissues have demonstrated that vibroacoustic signals encode major anatomical events such as entry into the peritoneal cavity in pigs [13] where the sudden change in mechanical resistance upon puncture led to a distinct vibroacoustic event. These studies established the fundamental feasibility of vibroacoustic sensing, but they primarily stem from simplified material systems and the experiments were designed to create a strong signal response (e.g. cavity entry) [11], [13]. In realistic biological tissue, which is characterized by contextualized tissue interconnected by fascia, surrounded by other soft-tissue structures and exposed to complex biomechanical constraints, signal interpretation becomes significantly more challenging. In such settings, vibroacoustic signals are more likely to exhibit a high density of events arising from needle bending due to insertion force, deformation, and other non-specific interactions. As a result, signal features cannot be unambiguously attributed to anatomical events based only on the vibroacoustic signal. Due to that, before moving vibroacoustic sensing into more advanced *in vivo* settings, a fundamental challenge remains: The lack of a reliable reference that enables the systematic investigation of how recorded signals relate to underlying biomechanical events during needle insertion. Solving this ambiguity requires an independent spatial reference that enables the targeted investigation of candidate events.

Post-procedural trajectory reconstruction represents a potential approach to provide this spatial reference. In this context, this methodological feasibility study proposes a spatiotemporal fusion pipeline that aligns soft-tissue crossings identified from post-procedure high-resolution micro-computed tomography (µCT) with vibroacoustic signals obtained during puncture. By reconstructing the needle trajectory and identifying tissue layer crossings in µCT, the expected number and spatial relationship of anatomical events can be approximated. Aligning these spatial annotations onto the signal domain by using the net displacement of the needle over time, enables to infer temporally constrained candidate regions for potential tissue interactions, and supports the interpretation of complex signal patterns.

The proposed pipeline was demonstrated with 22 annotated soft-tissue crossings across three different punctures, showcasing the technical implementation and applicability of aligning vibroacoustic signals with independently verified anatomical transitions. Instead of directly validating or detecting tissue interactions, this study provides a framework for constraining and systematically investigating candidate events in complex and high-density vibroacoustic signals. This methodological proof-of-concept is an indispensable prerequisite for future studies involving larger datasets and quantitative analysis for the development of automated detection of tissue transitions in vibroacoustic signals and the subsequent translation to more complex *ex vivo* and *in vivo* models.

## II. Methods

To explore whether post-procedural trajectory reconstruction can support the interpretation of ambiguous vibroacoustic signals, an experimental framework was developed aimed to provide (i) a biologically relevant soft-tissue model and (ii) an independent high-resolution reference modality to approximate internal tissue-layer crossings.

The experiments involved puncturing a soft-tissue model three times while simultaneously recording vibroacoustic signals and tracking needle velocity. Three different strategies were tested for visualizing the needle paths post-puncture to determine the most reliable method for trajectory reconstruction. The best marking strategy was used to annotate soft-tissue transitions along the needle trajectory and temporally align these spatial events with the recorded vibroacoustic signals.

### Model selection

The tobacco hornworm larva *Manduca sexta* (*M. sexta*) was selected as a biologically relevant model because its internal anatomy comprises multiple well-defined, interconnected soft-tissue layers and, unlike larger vertebrate animal models or *ex vivo* tissue, its size and geometry allow complete, high-resolution post-procedural micro-computed tomography (µCT) images [14]. By allowing visualization of the entire puncture trajectory, *M. sexta* uniquely provides a spatial reference for needle–tissue transitions on the micrometer scale, without relying on real-time imaging modalities that lack sufficient spatial resolution (Figure 1 A). Conventional phantoms fail to capture the complexity of biological tissue and bigger *ex vivo* models typically do not allow direct observation or precise reconstruction of internal needle trajectories. Intraoperative imaging modalities, such as ultrasound (US), lack the spatial resolution to resolve micro-scale transitions [1] and, consequently, currently there is no method capable of determining, with micrometer-scale accuracy, whether a needle has intersected or missed a specific internal tissue layer during puncture that should or should not have resulted in a vibroacoustic event. In this context, this model is used to develop and evaluate a spatiotemporal alignment framework that links vibroacoustic recordings to verified anatomical transitions, rather than to characterize native tissue mechanics or to directly translate to clinical tissue properties.

**Fig. 1.**
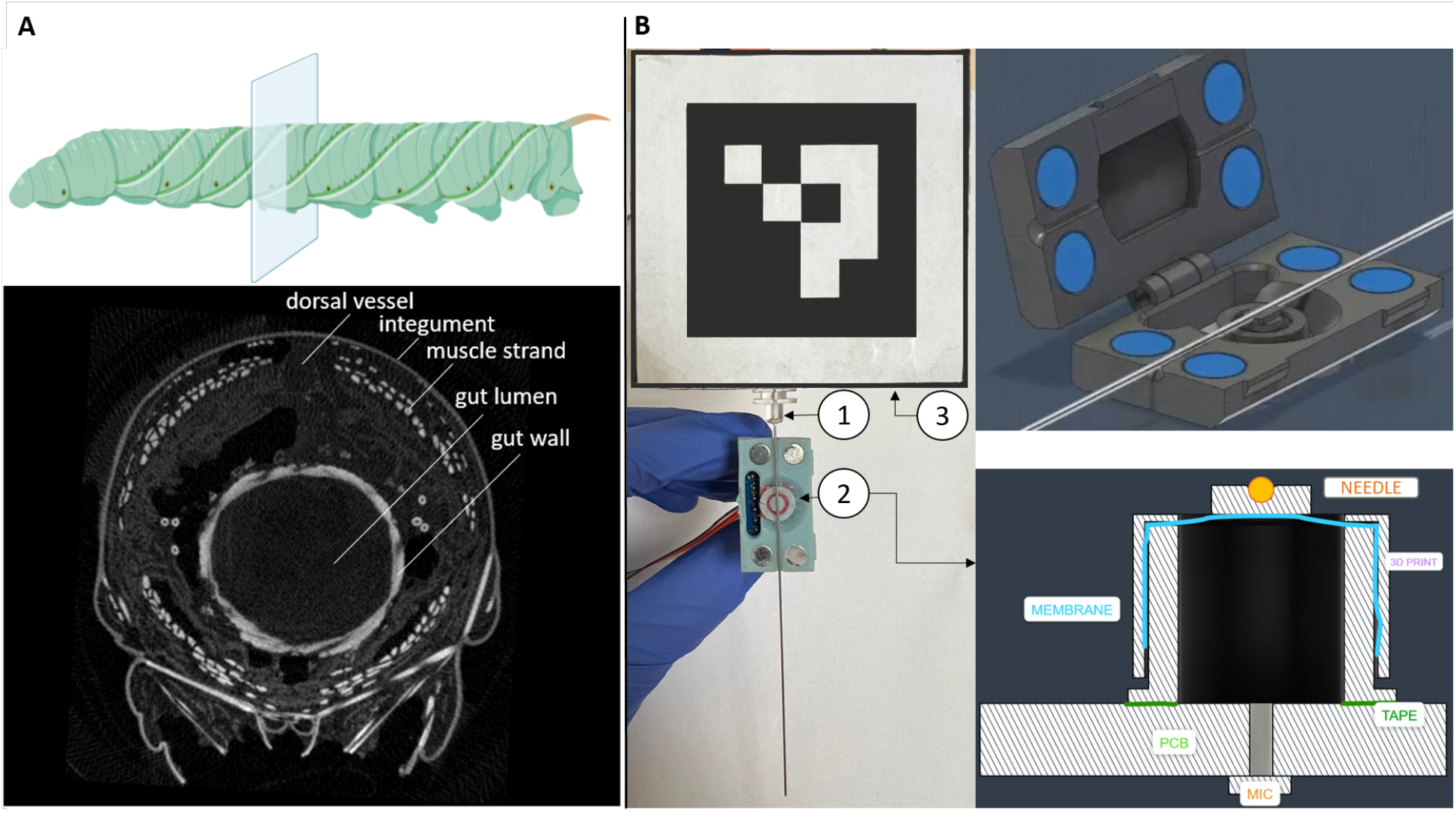
2D axial slice of M. sexta (left) and puncture setup (right) A) Important soft-tissue layers are marked. The gut lumen is filled with food mousse. The dorsal vessel acts as the insect equivalent of a pumping heart. Most muscle strands are arranged radially symmetrical and run horizontally. Created in BioRender [15] B) Left: close-up of a 24G needle (1) with one half of the vibroacoustic signal acquisition clip-on prototype attached to the proximal end of the needle shaft (2). The shaft is placed in front of a membrane (red) that funnels vibroacoustic signals to the microphones sound port. Needle holder with ArUco marker for velocity tracking (3). Right: schematic visualization of sensor-needle coupling and sound amplification funnel.

### Vibroacoustic Prototype and Signal Acquisition

To capture FIV generated during needle insertion a custom-designed vibroacoustic prototype was developed. It incorporates a Micro-Electro-Mechanical Systems (MEMS) microphone (Adafruit I2S MEMS Microphone Breakout, SPH0645LM4H) mounted on a three-dimensional (3D) printed clip-on module that is designed to attach the sensor to the proximal end of the shaft of standard medical needles (Figure 1 B top). The microphone was connected to a Raspberry Pi 4B that recorded vibroacoustic signals at 48 kHz via a custom laboratory acquisition application. The vibroacoustic sensor prototype is a further development of the most promising one presented in Oran *et al*. [8]. Instead of being positioned directly in front of the microphone’s sound port, the needle shaft is now coupled to a membrane that transfers and amplifies the signals into an underlying funnel, which leads them to the sound port (Figure 1 B right bottom).

The entire puncture procedure was recorded with two video cameras (Jabra PanaCast 20, 30 fps) from different angles to facilitate subsequent annotations and needle tracking. The needle was held on a 3D-printed holder with an ArUco marker attached for velocity tracking [16]. ArUco markers allow estimation of displacement and orientation based on their defined size and provide accurate tracking within ±2 mm as long as the marker is clearly visible within the camera’s field of view. In this study, the marker’s displacement between consecutive frames was used to compute the insertion velocity. This approach tracks velocity rather than absolute needle tip position, providing a consistent measure of net displacement over time that can be used to align µCT-derived spatial information with the temporal axis of the recorded vibroacoustic signals. The velocity estimation depends on the 30 fps video frame rate, introducing a temporal granularity of 33 ms, which means that vibroacoustic events are expected to occur within a narrow temporal window around the estimated tissue crossing times. A custom-made laboratory application was used to synchronize video and audio recordings using an external trigger present in both recordings. After synchronization, needle entry and exit were manually annotated to provide reference time points. A comprehensive methodology pipeline diagram is shown in Figure **??**.

Three *M. sexta* specimens were incubated in 70 % ethanol for 14 days to preserve internal structures, stabilized on a fixed surface, and punctured manually three times with a 24 G Quincke bevel needle with the vibroacoustic prototype attached. To enable post-procedural reconstruction of the needle trajectory, different marking strategies were evaluated.

1. Pre-coating the needle with an iodine-based contrast agent (Gastrographin) or injecting contrast agent upon needle retraction to highlight the puncture path (n=3).
2. Detecting a “negative trajectory” by analyzing areas of tissue disruption caused by puncture (n=3).
3. Inserting a µCT-compatible nylon string (0.21 mm) (n=3) or tungsten wire (0.10–0.21 mm)(n=3) through the needle before withdrawal to preserve the path.

After the trajectory was marked, the specimen was incubated in 4 % Molybdatophosphoric acid hydrate (PMA) dissolved in 70 % ethanol for 14 days to enhance soft-tissue contrast, before imaging with high-resolution µCT.

### µCT Imaging, Image Processing and Multimodal Fusion

The soft-tissue layers in *M. sexta* are only 0.1 to 0.2 mm thick, requiring high resolution to determine the crossing point of tissue puncture. µCT imaging features those resolutions down to 4 µm and was consequently chosen as the reference modality for this study (SkyScan 1173, Bruker µCT, Belgium at 50 kV, 160 µA, 950 ms exposure, and 900 projections over 360°, 24 µm isotropic voxel size, total scan time of 40–50 minutes per specimen). Reconstruction was performed using NRecon GPUReconServer (v2.1.0) with Hamming filtering and ring-artifact correction level 7. The resulting 16-bit BMP image stacks were processed in Amira 3D Pro v. 2022.1 (ThermoFisher Scientific, Massachusetts) to visualize anatomy, segment trajectories, and export spatial coordinates of soft-tissue layer crossings along the trajectories as .csv files. For this study the entire trajectories were segmented and visualized as a 3D rendering to illustrate the relationship between the puncture path and the anatomical layers. For future high-throughput analysis and automation, 2D axial slices will be sufficient to annotate tissue boundaries and correlate them with vibroacoustic events.

A multimodal fusion pipeline was developed that combined the needle velocity obtained from video using ArUco marker detection, visual annotation of needle entry, the coordinates of soft-tissue layer crossings derived from the µCT images, and the recorded vibroacoustic waveform.

The needle velocity data were smoothed using a three-frame rolling mean. Inside the manually annotated entry event, the alignment was started immediately after the rupture of the integument, defined by a high energy peak inside the vibroacoustic signal and a sharp downward movement in the velocity plot. This point was set to zero in the plots to allow the observation of net displacement over the puncture. Based on the relative distances between µCT-annotated tissue crossings and the integrated velocity to derive the cumulative insertion depth over time, the expected frame indices of each layer crossing were calculated and mapped to the corresponding audio timestamps.

For visualization, the vibroacoustic signal’s amplitude envelope was plotted with color-coded markers indicating aligned tissue crossings within a 300 ms window, along a synchronized Continuous Wavelet Transform (CWT) spectrum and a plot showing the net displacement in mm. Vibroacoustic events are not expected to occur at these exact time points but rather within a temporal neighborhood of ±150 ms (300 ms total) around each calculated tissue crossing. This choice reflects the physical duration of FIV events, which typically extend beyond 300 ms for prominent interactions such as cavity entry [13], with smaller-scale transitions expected to be shorter yet still distributed in time [10]. Although tissue crossings are determined with micrometer-scale precision from µCT annotations, mapping these spatial positions to the time axis requires identifying the exact moment of tissue rupture as a registration reference. At 30 fps video frame rate and a sampling rate of 48 kHz, a small temporal uncertainty of 33 ms (1600 samples) is introduced at choosing the starting point for the alignment. At the observed mean velocities of 1-2 mm/s, 33 ms corresponds to a spatial offset of up to 60 µm. Therefore, the window reflects the physically distributed nature of FIV and possible registration inaccuracies.

## III. Results

Three trajectory marking methods were tested to determine a suitable approach for reconstructing needle trajectories in soft tissue. The selected method was subsequently used as the basis for marking soft-tissue crossings for subsequent mapping of vibroacoustic signal events to anatomical transitions.

### Evaluation of trajectory marking strategies

Injecting contrast agent upon needle retraction or pre-coating the needle with contrast agent before puncture did not yield visible trajectories as the contrast diffused into surrounding tissue, preventing the visualization of a defined puncture path (Figure2 B). The use of metallic tungsten wires (0.25 mm and 0.1 mm) enabled partial visualization of the trajectory, but caused substantial X-ray artifacts that obscured the surrounding anatomy (Figure2 A). Both wires deformed during insertion, distorting the true puncture trajectory. The thicker wire tended to spring back to its original form, while the thinner wire bent inside the tissue. This approach also required two separate µCT scans per specimen to allow subsequent overlapping of anatomical structures, which is unsuitable for high-throughput studies, increases acquisition time and risk of reconstruction errors. Therefore, we concluded that tungsten wire is unsuitable as marker for fine-scale trajectory preservation in soft-tissue models.

**Fig. 2.**
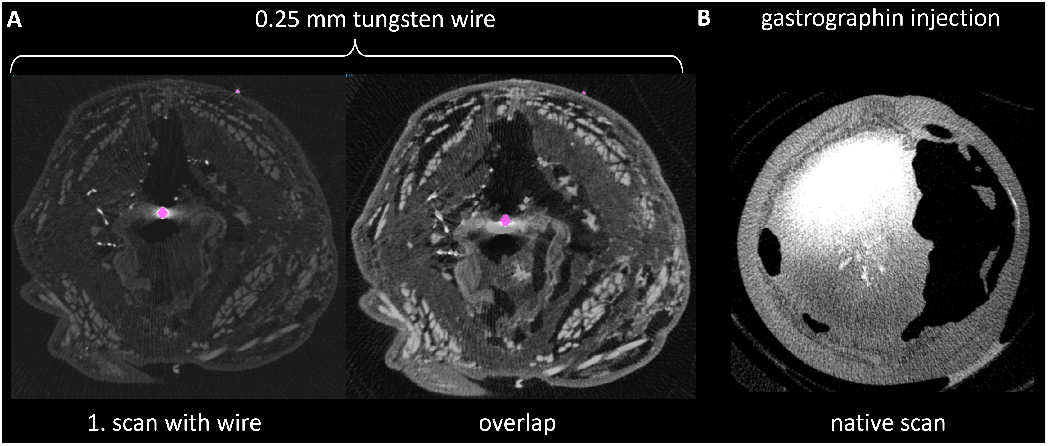
Axial slices of M. sexta specimen with tungsten wires and contrast agent for trajectory marking. A: The tungsten wire (pink) introduces geometric deformation and X-ray artifacts in the 1. scan. A second scan was run after removing the wire and registered with the first scan to allow for a more clear visualization of the anatomy. Two independent scans are resource heavy and introduce the risk of registration error, making tungsten wires unsuitable for trajectory reconstruction. B: Upon needle retraction, gastrographin contrast agent was injected to mark the trajectory. In the native state scan no distinct trajectory was observable because the contrast agent bled into adjacent tissues.

A µCT-compatible nylon string (0.21 mm) yielded the best results, by introducing minimal artifacts and maintaining stable geometry without altering anatomy (Figure 3). This approach enabled simultaneous visualization of anatomical context and trajectory within a single µCT scan per specimen and was therefore selected for subsequent multimodal fusion.

**Fig. 3.**
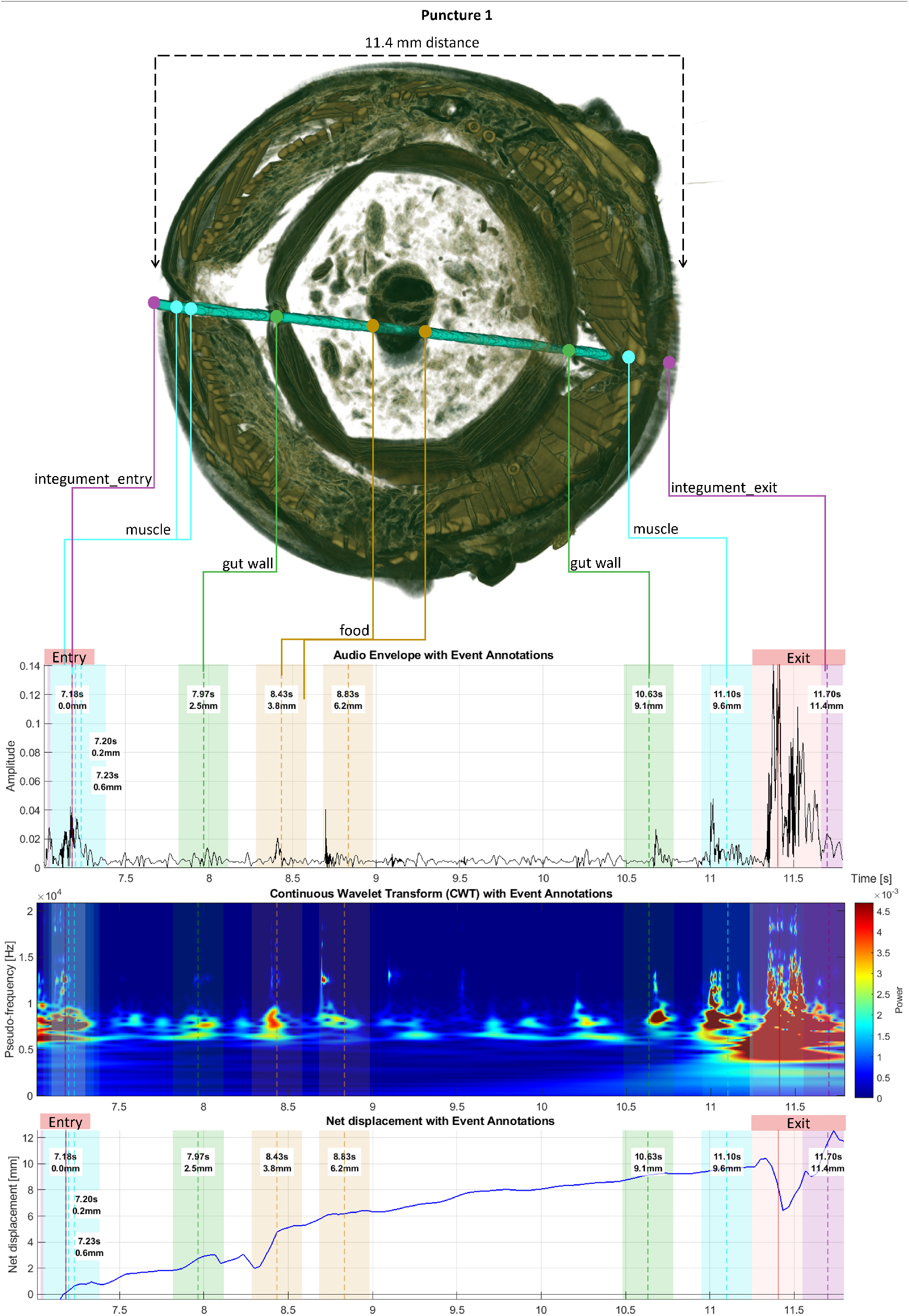
Spatiotemporal alignment of µCT-based trajectory reconstruction and vibroacoustic signals. Top: Axial view of a 3D reconstruction of a M. sexta specimen with a 0.25 mm nylon string segmented (cyan) and annotated soft-tissue transitions. Bottom: corresponding vibroacoustic envelope, CWT spectrogram, and net needle displacement during puncture. µCT annotated tissue transitions were mapped to the vibroacoustic envelope and CWT spectrum using the needle net displacement. Calculated hit times are shown within a 300 ms window. Peaks in the vibroacoustic envelope and events in the CWT spectrum align with entry and exit punctures, as well as additional transitions potentially corresponding to muscle, gut-wall and food crossings. The moment after integument rupture during the entry event (red line) was used as starting point for temporal alignment.

### Feasibility of Spatiotemporal Fusion

For the spatiotemporal alignment each crossed anatomical layer along the segmented trajectory was mapped to the vibroacoustic waveform and CWT spectrogram. The temporal appearance of each soft-tissue transitions in the vibroacoustic events was determined computationally by combining the net displacement over time with the spatial positions of layers from the µCT-based 3D reconstruction (Figure 3). The alignment was initialized using the rupture of the integument as starting reference. For all punctures, the calculated end of the trajectory corresponded closely to the observed exit event in the vibroacoustic signal, indicating that the velocity data provided a plausible approximation of needle progression over time.

The spatiotemporal fusion was tested with 22 annotated layer crossings from three different punctures, resulting in similar observations (Figure 3 and 4), indicating that the fusion pipeline can be consistently applied across repeated insertions and yields plausible temporal relationships. Distinct vibroacoustic events appeared within a defined temporal neighborhood of the calculated tissue crossings rather than at single discrete instants. Entry and exit punctures were confirmed by manual annotations. The mean velocity throughout all punctures was 1-2 mm/s. Especially upon crossing the integument (purple window at 7.18 s and 11.70 s), a high-amplitude peak with low-frequency components was observed. Smaller events were observed in temporal proximity to gut-wall and muscle crossings. Inside the gut, food particles were visible in the µCT reconstruction, and vibroacoustic events were observed in temporal proximity to these structures.

**Fig. 4.**
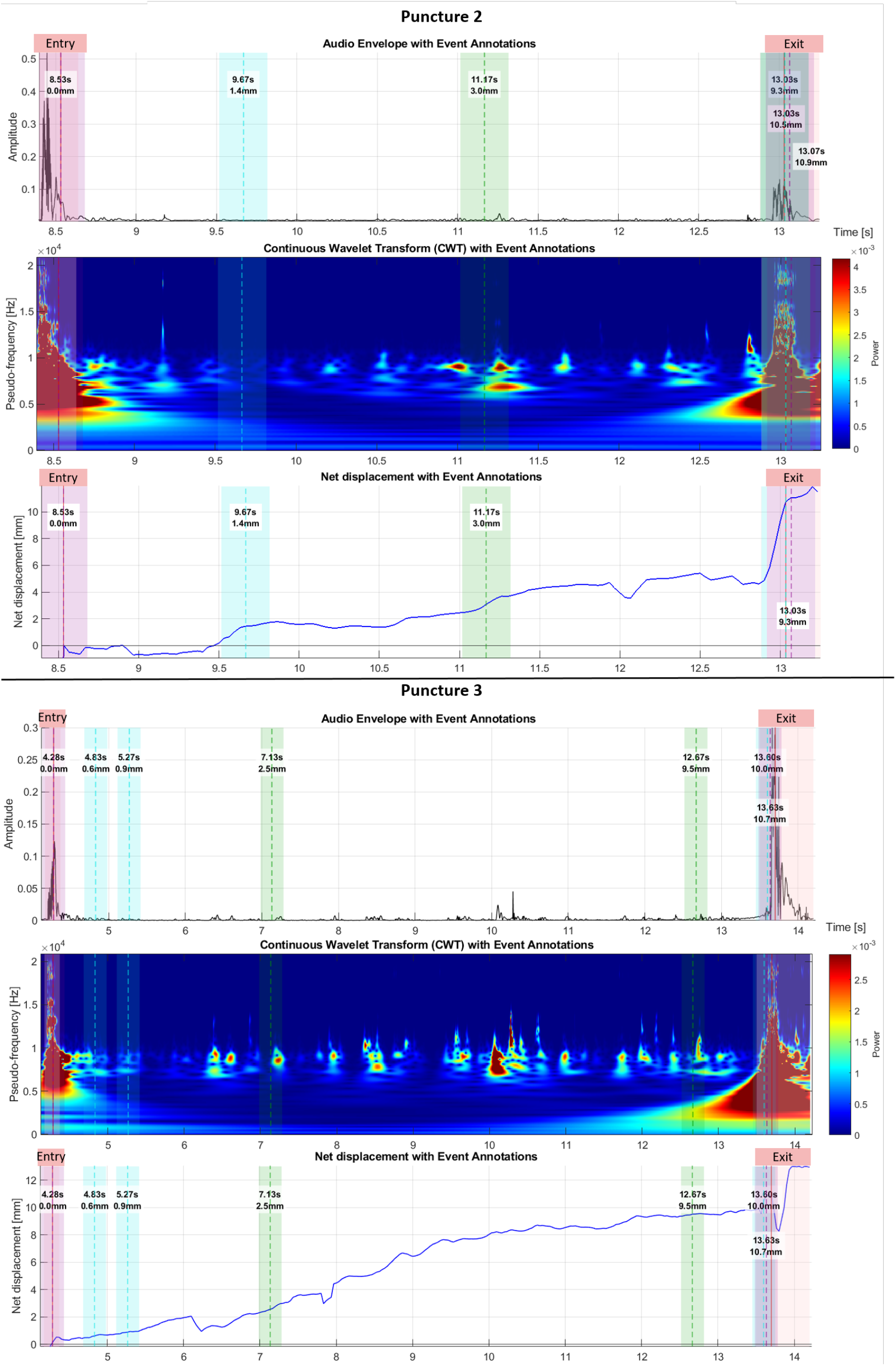
Spatiotemporal alignment of puncture 2 and 3. purple: integument crossing; blue: muscle crossing; green: gut wall crossing; red: manually annotated entry and exit punctures. Events occurring between the two gut wall crossings correspond to food particles within the gut (not annotated), consistent with the 3D reconstruction shown in Figure 4. The point of maximum energy during the entry event (red line) was used as a reference point for temporal alignment.

Manual insertions enabled force modulation as a response to transient resistance during tissue penetration, which was reflected in the net displacement profiles. For example, in Puncture 1, distinct reductions in net displacement were observed upon crossing a food particle at 8.43 s and during exit at 11.4 s, followed by immediate recovery once the resistance was overcome.

The detailed reconstruction of puncture trajectories provided important information for interpreting the vibroacoustic signals at that state, that would not be accessible using other modalities (Figure 5). For example, contrary to initial expectations, in both dorsal punctures (puncture 1 and 2), the dorsal vessel was not intersected due to tissue deformation, indicating that a corresponding vibroacoustic event should not be expected from this layer (Figure 5 A). Additionally, for puncture 1, it was observed that the needle passed through a food clump within the gut lumen. This observation allowed to annotate this as additional candidate events, which helped finding a plausable connection for the vibroacoustic events at 8.43 s and 8.83 s (see figure 3) ref. Figure 5 B). Furthermore, two trajectories from separate punctures were observed in close proximity to each other, where the 3D visualization clarified that those two did not overlap (Figure 5 C).

**Fig. 5.**
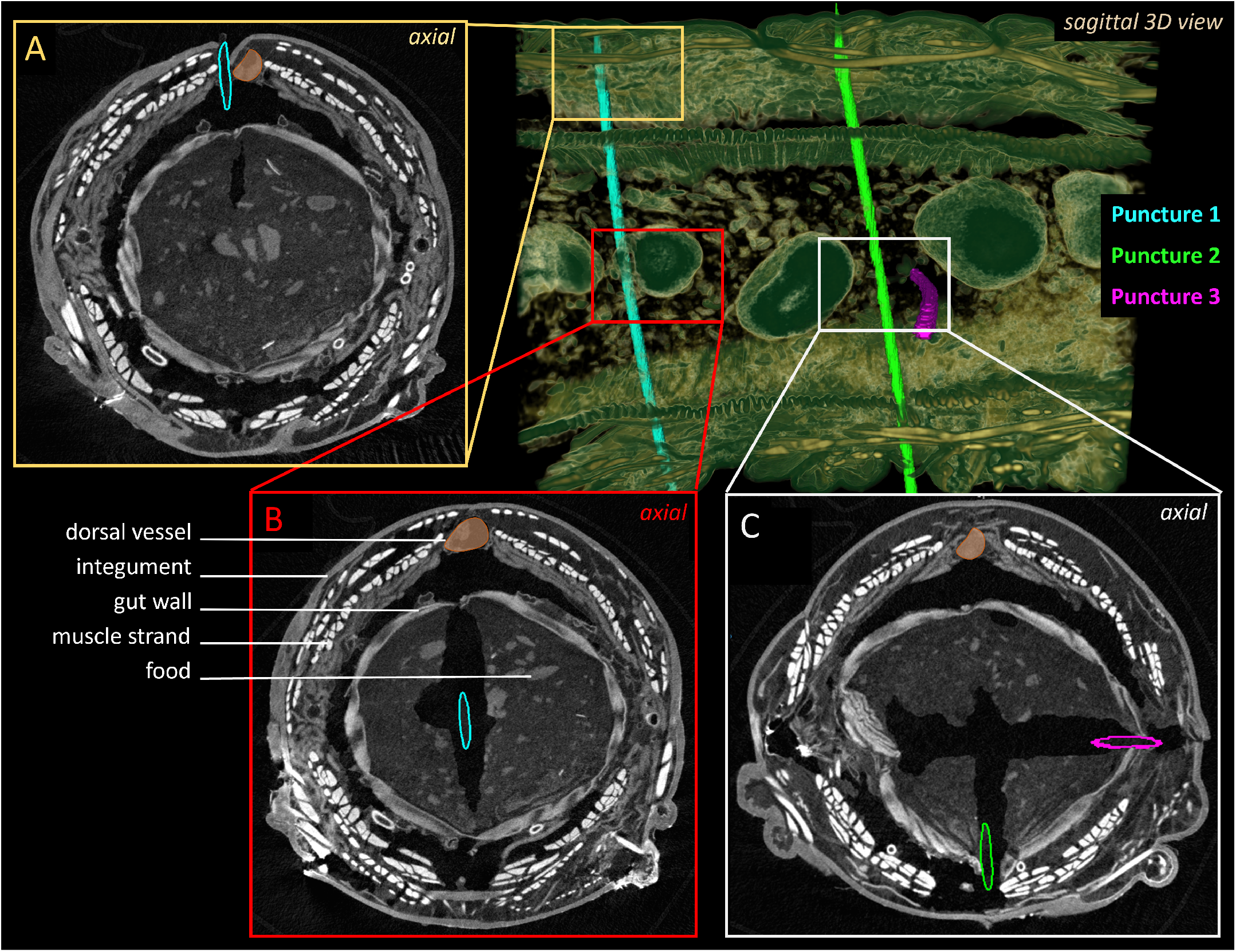
µCT enables observation of needle trajectories in high-resolution and allows to align anatomical events with vibroacoustic signals. Saggital 3D view of the punctured M. sexta specimen with 3 segmented trajectories as examples and axial close up views. A) During puncture 1 the dorsal vessel was missed due to dislocation. Knowing this, no vibroacoustic event was expected in the temporal alignment of that event. B) The needle clearly punctured a clump of food mousse inside the gut. Entry and exit into the food ball were annotated and mapped to the vibroacoustic signal (see figure 3). C) Two punctures were close to each other. The needle removed surrounding tissue but the exact trajectories, visualized with the nylon strings, did not cross. Additionally it can be observed that the punctures were not executed in 90°angle but tilted a little from top left to bottom right. Those observations would not be possible without the µCT images.

**Fig. 6.**
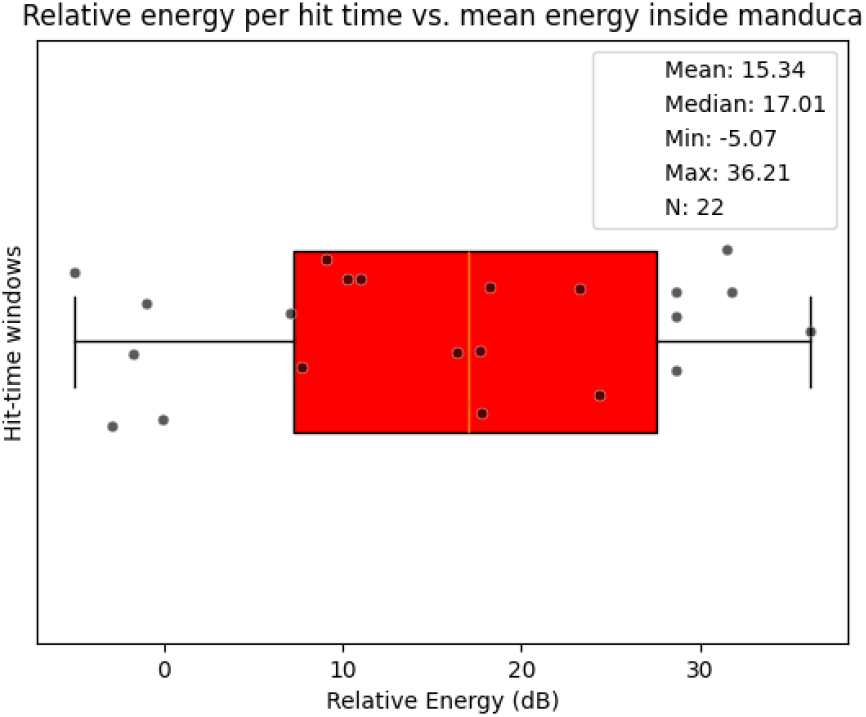
The energy inside the windows with expected signal crossings exceeded the mean energy inside the larva in 77,3 % of the cases. The energy inside 22 aligned windows was compared to the mean energy inside the larva during this puncture. 17 out of 22 potential tissue crossings fell in the temporal proximity of high-energy signal regions.

These observations demonstrate the value of µCT as an independent spatial reference for initially interpreting vibroacoustic signals, particularly in complex environments where multiple plausible signal sources exist.

To assess whether the aligned windows tended to coincide with regions of increased signal activity, the mean energy within the defined temporal windows around the mapped crossing times was compared to the mean energy of the signal within the larva (Figure.

In 17 out of 22 cases (77.3 %), the energy within the aligned windows exceeded the average signal energy. However, obviously this effect depends on the overall complexity of the signal. In signals with lower event density, the higher relative energy within aligned windows suggests that the alignment tends to capture regions of increased activity rather than ending up in uniform signal regions. In punctures with a high density of events (e.g. puncture 3), the relative energy within aligned windows does not exceed the global mean, reflecting the elevated baseline activity.

## IV. Discussion

This study established and demonstrated a multimodal pipeline for qualitatively mapping vibroacoustic events with anatomical transitions. By aligning trajectory reconstructions from µCT-images with vibroacoustic signals, the pipeline provides an anatomically grounded reference for interpreting acoustic events in relation to soft-tissue interactions. This fills a methodological gap in vibroacoustic research, where signal events were difficult to interpret or risked being confounded by artifacts because of the lack of a biological reference.

Previous studies in vibroacoustic sensing primarily relied on artificial phantoms or *ex vivo* tissues embedded in gelatine, which enabled controlled experiments but lacked the complexity of interconnected systems [7], [10], [11]. Work on pig cadavers demonstrated the detection of major puncture events, such as cavity entry, but finer soft-tissue transitions remained unresolved because of the lack of ground truth [13]. *M. sexta* used in our study, represents a logical next step between artificial and mammalian models because it features multiple interconnected soft-tissue layers while remaining small enough for complete visualization using µCT. Using ethanol-preserved specimens helped minimize tissue deformation during puncture, which was essential for the study’s goal of demonstrating the proposed spatiotemporal fusion methodology in a controlled environment. Therefore, the results do not permit direct translation of tissue-specific vibroacoustic characteristics into real clinical applications, but represent a necessary prerequisite for future studies in less observable *ex vivo* and *in vivo* systems.

During needle insertion, complex biomechanical interactions between the needle and the tissue generate forces that can be modeled mathematically and physically. These models incorporate tissue contact stiffness, friction along the needle shaft, and cutting forces at the tip, but do not capture the full complexity of living tissue [4], [17]–[19]. These biomechanical forces are challenging to simulate but might be encoded in vibroacoustic signals, potentially providing a measurable derivation of tissue interaction. For example, when a tissue barrier ruptures, force accumulates and creates a characteristic vibration pattern upon release. This sharp amplitude peak, followed by rapid decay within milliseconds, was characterized as the moment of cavity puncture [13]. Similarly, preliminary observations from three *M. sexta* punctures showed lower-energy peaks of a qualitatively similar shape during integument puncture. Events in timely proximity to muscle crossings generally produced the lowest-intensity signals and occasionally overlapped with entry or exit events. In puncture 2, one muscle strand was either displaced during insertion or did not generate a discernible vibroacoustic event. This could be explained by the predominantly horizontal and radially symmetric arrangement of muscle strands in *M. sexta* (Figure 1A), which can cause them to be shifted during puncture. This observation underscores the value of combining vibroacoustic sensing with µCT-based reconstruction to draw meaningful conclusions about actual needle–tissue interactions. Current robotic-assisted needle procedures are limited by preoperative imaging that cannot account for tissue deformation and by intraoperative ultrasound with insufficient resolution and absent haptic feedback [2], [20], [21]. These challenges highlighted the need for a feedback channel, like vibroacoustic sensing, that provides real-time information on tissue integrity and structural transitions as additional navigation information.

Before this can be achieved, several limitations remain. The accuracy of the multimodal alignment is limited by the 30 fps video frame rate, which, together with a sample rate of 48 kHz, introduces a temporal uncertainty of 33 ms (1600 samples). At the observed mean velocities of 1-2 mm/s, this corresponds to a spatial offset of up to 60 µm. Since tracking is used only to align µCT-derived spatial information with the temporal axis of net needle displacement, and not to determine the absolute needle tip position, this uncertainty limits point-wise alignment but does not invalidate the assessment of temporal proximity between vibroacoustic events and underlying tissue interactions. However, at higher insertion velocities, this offset will increase proportionally, leading to larger alignment errors. As a result, higher frame rates will be required for faster insertions.

This study focused on establishing and demonstrating the technical feasibility of the proposed multimodal fusion pipeline. The objective was not to assess biological variability or generalize vibroacoustic signal characteristics across species, but to determine whether the data acquisition pipeline is suitable to spatiotemporally align vibroacoustic signals with µCT-derived anatomical information. Using a small number of punctures in a structurally interconnected soft-tissue model provided a controlled environment to evaluate this fundamental methodological capability question before introducing additional biological variability.

## V. Conclusion

This study presents a multimodal framework that bridges high-resolution anatomical imaging and vibroacoustic sensing to investigate needle–tissue interactions at a very high level of detail. By combining µCT-based trajectory reconstruction with synchronized vibroacoustic recordings, we established a biologically grounded reference for interpreting acoustic events in relation to verified tissue transitions. The proposed methodology enables the systematic exploration of how mechanical interactions within complex soft-tissue environments are reflected in vibroacoustic signatures, addressing a critical challenge that has limited progress in the field.

Beyond demonstrating technical feasibility, this work introduces a new paradigm for studying procedural sensing. Rather than relying solely on force models or imaging feedback, vibroacoustic sensing captures the dynamic mechanical consequences of tissue interaction directly at the instrument–tissue interface. The presented framework provides the foundation required to build annotated datasets, develop quantitative biomechanical models, and train machine-learning algorithms capable of recognizing tissue transitions and procedural events automatically.

In the long term, vibroacoustic sensing could evolve into a complementary feedback channel for image-guided and robotic interventions, enabling real-time detection of tissue boundaries, characterization of tissue integrity, and adaptive trajectory control. Such capabilities may help compensate for the loss of tactile perception in robotic systems and provide information at temporal and spatial resolutions beyond those achievable with conventional intraoperative imaging.

Ultimately, the convergence of vibroacoustic sensing, computational modeling, and artificial intelligence may enable future needles and catheters to become self-aware sensing instruments that continuously interpret their surrounding tissue environment, contributing to safer, more precise, and increasingly autonomous minimally invasive interventions.

## Acknowledgment

The authors thank Jessica Steinbart for her insight into µCT imaging.

## Data availability

The data is available on Huggingface: https://huggingface.co/datasets/VibroNav/ManducaEventFinder112025. For issues related to the data files you may contact K.Steeg.

## Ethical statement

The animals used for this study are not subject to ethical constraints.

